# How Myopia and Glaucoma Influence the Biomechanical Susceptibility of the Optic Nerve Head

**DOI:** 10.1101/2022.12.19.520997

**Authors:** Thanadet Chuangsuwanich, Tin A. Tun, Fabian A. Braeu, Clarice H.Y. Yeoh, Rachel S. Chong, Xiaofei Wang, Tin Aung, Quan V. Hoang, Michaël J.A. Girard

## Abstract

**Purpose:** We aimed to assess optic nerve head (ONH) deformations following acute intraocular pressure (IOP) elevations and horizontal eye movements (adduction and abduction) in control eyes, highly myopic (HM) eyes, HM eyes with glaucoma (HMG), and eyes with pathologic myopia alone (PM) or PM with staphyloma (PM+S).

**Methods:** We studied 282 eyes, comprising of 99 controls, 51 HM, 35 HMG, 21 PM and 75 PM+S eyes. For each eye, we imaged the ONH using spectral-domain optical coherence tomography (OCT) under the following conditions: **(1)** primary gaze, **(2)** 20° adduction, **(3)** 20° abduction and **(4)** primary gaze with acute IOP elevation (to ~35 mmHg) achieved through ophthalmodynamometry. For each OCT volume, we automatically segmented the ONH tissues using deep learning. We performed digital volume correlation (DVC) analysis to compute IOP- and gaze-induced ONH displacements and effective strains (i.e. local deformations). All biomechanical quantities were compared across groups.

**Results:** Under IOP elevation, we found that HM eyes exhibited significantly lower strains (3.9 ± 2.4 %) than PM eyes (6.9 ± 5.0%, p < 0.001), HMG eyes (4.7 ± 1.8%, p = 0.04) and PM+S eyes (7.0 ± 5.2%, p < 0.001). Under adduction, we found that HM eyes exhibited significantly lower strains (4.8% ± 2.7%) than PM+S eyes (6.0 ± 3.1%, p = 0.02). We also found significant associations between axial length (or refractive error) and strains - eyes with higher axial length and greater myopia were associated with higher strains. IOP-induced strains were also positively correlated with adduction-induced strains.

**Conclusion:** We found that HMG eyes experienced significantly higher strains under IOP elevations as compared to HM eyes. Additionally, PM+S eyes experienced highest ONH strains as compared to other groups under all biomechanical loads. Our preliminary findings suggest the possibility of using a simple biomechanical test to tease out the susceptibility of HM eyes to further develop glaucoma and/or staphyloma.

## Introduction

An association between glaucoma and myopia is well known and it has been reported in many clinical and population-based studies.^1–3^ Numerous studies have suggested that moderate to high myopia is associated with increased risk of primary open angle glaucoma, normal-tension glaucoma, and ocular hypertension.^4–7^ Myopic individuals have also been found to be more at risk of having an elevated intraocular pressure (IOP) than the general population.^8–10^ However, to date, reports on the associations between myopia and glaucoma are largely observational^11^ and the pathological link between the two conditions is still speculative.

According to the biomechanical theory of glaucoma pathogenesis, it is reasonable that myopia is a risk factor for glaucoma, since many biomechanical changes in myopia lead to structural weakening of the optic nerve head (ONH).^12^ In myopia, the eye elongates, and the posterior sclera remodels, thins and weakens, exhibiting a general loss of collagen and proteoglycans.^13, 14^ The scleral collagen fibers become thinner, and reorganize into a lamellar arrangement (rather than interwoven).^15^ In addition, the lamina cribrosa thins and the sclera canal enlarges.^16, 17^ The sclera also exhibits an increased creep rate (a viscoelastic property) indicating facilitated sliding of scleral lamellae.^18^ Although a signalling pathway between the retina and the posterior sclera has been suggested,^19^ it remains unclear why these biomechanical changes occur in myopia. Overall, these changes could make the ONH more susceptible to biomechanical forces exerted by IOP, causing damage to the axons at the lamina cribrosa (LC) and leading to glaucoma development.^17^

Several studies have employed biomechanical stress tests to assess the robustness of the ONH, using either IOP, or optic nerve traction as the primary mechanical stimulus (ours and others).^20–25^ Previous literatures have used such tests to elucidate the role of IOP elevation or optic nerve traction in glaucoma pathogenesis, but no studies have investigated the in vivo ONH responses of a spectrum of myopic eyes under biomechanical loads. With much evidence suggesting a possible biomechanical link between glaucoma and myopia, we believe that a biomechanical stress test to tease out ONH structural robustness in myopic eyes will give us further insight into the link between the two diseases; this may also help us identify myopic individuals that are at risk of developing glaucoma.

In addition, providing an accurate glaucoma diagnosis in a highly myopic patient remains a major clinical challenge. This is because conventional biomarkers for glaucoma diagnosis (e.g. ONH appearance, peripapillary atrophy and visual field defect on perimetry)^26, 27^ are greatly obscured by the structural alterations typically found in high myopia (HM) (e.g. myopic peripapillary atrophy, tilted disc and retinal nerve fiber layer [RNFL] thinning)^26, 28^ and pathologic myopia (PM) (e.g. geographic atrophic changes in myopic macular degeneration that result in visual field defects).^11, 29, 30^ With these conventional limitations, a biomechanical stress test has the potential to fill in this gap in identifying whether a given eye is glaucomatous, myopic, or a combination of both.

In this study we aimed to assess the biomechanics of the ONH in patients with a different spectrum of myopia-related conditions, which include HM, PM and pathologic myopia with staphyloma (PM+S), and in those that were confirmed to exhibit both myopia and glaucoma (HMG). Our long-term goal is to characterize myopia-related conditions in terms of their biomechanical sensitivity to external loads and to tease out the possibility of using a biomechanical test to identify HM individuals who are at risk of developing glaucoma.

## Methods

### Subjects Recruitment

We studied 282 eyes, which comprised of 99 controls, 51 HM, 35 HMG, 21 PM and 75 PM+S from glaucoma clinics at the Singapore National Eye Centre. We included subjects aged more than 50 years old, of Chinese ethnicity (predominant in Singapore) and excluded subjects who underwent prior intraocular/orbital/brain surgeries, subjects with past history of strabismus, ocular trauma, ocular motor palsies, orbital/brain tumors; with clinically abnormal saccadic or pursuit eye movements; subjects with poor LC visibility in ocular coherence tomography (OCT) images (<50% en-face visibility); subjects with known carotid or peripheral vascular disease; or with any other abnormal ocular conditions.

In this study, control eyes had refractive error between +2.75 and −2.75 diopters with axial lengths less than 25 mm. Control eyes had normal IOP and no pathological features on the ONH and the retina. HM was defined by eyes with axial lengths greater than 25 mm and myopic refractive errors worse than 5 diopters. HMG was defined for eyes with both HM (excluding PM and staphyloma) and a glaucoma diagnosis as performed by a glaucoma specialist (RSC, AT). PM (without staphyloma) was defined for eyes exhibiting both HM and myopic macular degeneration as defined by the meta-analysis for PM classification system,^31^ or myopic traction maculopathy (without the presence of staphyloma). PM+S was defined as HM eyes with a clinical diagnosis of staphyloma based on multimodal imaging (fundus photography, optical coherence tomography, ultrasonography and in selected subjects, magnetic resonance imaging) and confirmed by a retinal specialist.

Glaucoma was defined as glaucomatous optic neuropathy (based on Nagaoka et al, 2015),^32^ characterized as loss of neuroretinal rim with vertical cup-to-disc ratio >0.7 or focal notching with nerve fiber layer defect attributable to glaucoma (based on a clinical observation) and/or asymmetry of cup-to disc ratio between eyes >0.2, with repeatable glaucomatous visual field defects (independent of the IOP value) in at least 1 eye. Glaucomatous visual field defect was defined the following conditions were present: (1) glaucoma hemifield test outside normal limits, (2) a cluster of ≥ 3, non-edge, contiguous points on the pattern deviation plot, not crossing the horizontal meridian with a probability of < 5% being present in age-matched normal (one of which was < 1%) and (3) Pattern Standard Deviation (PSD) < 0.05; these were repeatable on two separate occasions.

We included subjects aged more than 50 years old, of Chinese ethnicity (predominant in Singapore) and excluded subjects who underwent prior intraocular/orbital/brain surgeries, subjects with past history of strabismus, ocular trauma, ocular motor palsies, orbital/brain tumors; with clinically abnormal saccadic or pursuit eye movements; subjects with poor LC visibility in OCT (<50% en-face visibility); the method of determining LC visibility was given in our previous study.^33^ subjects with known carotid or peripheral vascular disease; or with any other abnormal ophthalmic and neurological conditions.

Each subject underwent the following ocular examinations: (1) measurement of refraction using an autokeratometer (RK-5; Canon, Tokyo, Japan) and (2) measurement of axial length, central corneal thickness and anterior chamber depth using a commercial device (Lenstar LS 900; Haag-Streit AG, Switzerland). For each tested eye we performed a visual field test using a standard achromatic perimetry with the Humphrey Field Analyser (Carl Zeiss Meditec, Dublin, CA).

This study was approved by the SingHealth Centralized Institutional Review Board and adhered to the tenets of the Declaration of Helsinki. Written informed consent was obtained from each subject.

### OCT Imaging

Subjects’ pupils were dilated with 1.0% Tropicamide before imaging with spectral-domain OCT (Spectralis; Heidelberg Engineering GmbH, Heidelberg, Germany). The imaging protocol was similar to that from our previous work.^34^ In brief, we conducted a raster scan of the ONH (covering a rectangular region of 15° × 10° centered at the ONH), comprising of 97 serial B-scans, with each B-scan comprising of 384 A-scans (**Figure 1a**). The average distance between B-scans was 35.1 μm and the axial and lateral B-scan pixel resolution were on average 3.87 μm and 11.5 μm respectively. All B-scans were averaged 20 times during acquisition to reduce speckle noise. Each eye was scanned four times under four different conditions – primary OCT position, 20° adduction, 20° abduction and acute IOP elevation.

**Figure 1.**
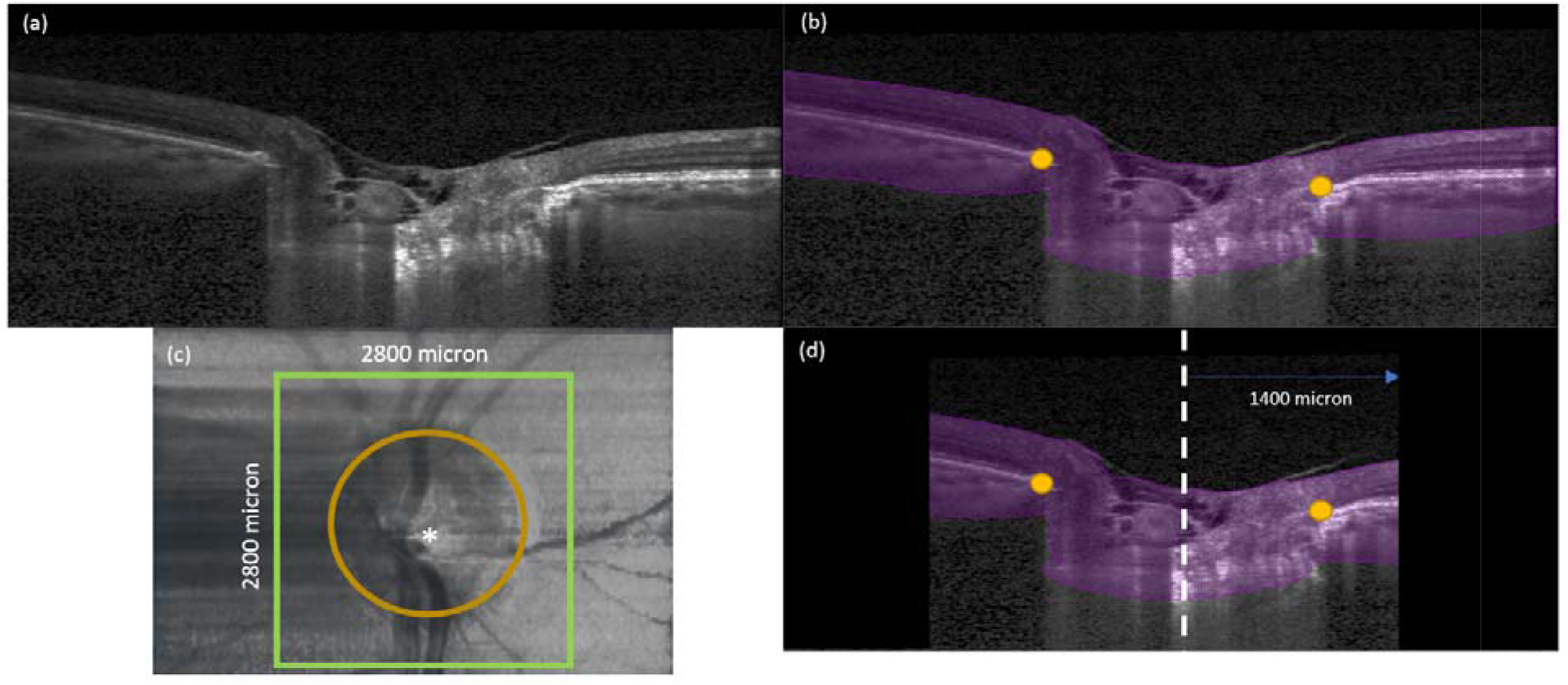
**(a)** A single B-scan obtained from the OCT machine without any image enhancement **(b)** Automatic segmentation of the B-scan in (a). ONH tissues were segmented in purple. In addition, BMOs (orange dots) were automatically marked for each B-scan **(c)** Anterior-surface view of the ONH. The ONH center (white star) was identified from the best-fit circle to the BMOs (orange-dotted line). Green square defines our region of interest to be cropped from the OCT volume with 2800μm length on each side. **(d)** A B-scan view after we apply cropping to the OCT volumes. Black region was not considered for our deformation tracking. The length from central line (white-dotted line) to the cropping border (green line) is 1400 μm.

### OCT imaging in primary gaze and in Adduction/Abduction positions

In this study, the primary gaze OCT position referred to the eye position during a standard OCT scan. This position does not exactly correspond to the primary gaze position as both the pupil and ONH need to be aligned with the OCT objective, inducing a slight eye rotation to the left in a right eye, and vice versa^34^. Amplitudes of horizontal gaze positions reported in this study were therefore with respect to the primary gaze OCT position. Procedures for imaging under different gaze positions have been described in our previous work^34^. Briefly, we employed a custom-built 3D printed rotatable chin rest to induce 20° adduction and 20° abduction and one OCT volume was acquired in each position.

### OCT imaging during acute IOP elevation

For each eye in the primary gaze position, we applied a constant force of 0.65 N to the temporal side of the lower eyelid using an opthalmodynamometer (ODM), as per an established protocol.^20, 21, 34^ The force was applied for approximately 2 to 3 minutes throughout the OCT scan duration and varied for no more than one minute across subject. This force raised IOP to about 35 mmHg and was maintained constant. IOP was then re-assessed with a Tono-Pen (Reichert Instruments GmbH, Munich, Germany), and the ONH was imaged with OCT in baseline position immediately (within 30 seconds) after the IOP was measured.

### ONH Tissues Segmentation

For each OCT volume, we automatically segmented the region contained by the ONH (**Figure 1b**) as defined by an anterior boundary (i.e. the inner limiting membrane) and a posterior boundary. The latter was represented by the posterior boundaries of the peripapillary sclera and of the LC, whenever visible. Otherwise, the visible portion of the OCT signal helped us define a posterior boundary. In contrast to our previous publications, no effort was made to isolate each neural and connective tissue layer at the ONH, as this task proved relatively difficult in myopic eyes with highly tilted ONH and pathological characteristics that obscured tissue boundaries. Segmentation was instead performed using a deep-learning algorithm similar to that designed in our previous work; it is a necessary step to perform 3D strain mapping.^35^ Bruch’s membrane opening (BMO) points were also automatically extracted with a custom algorithm. Note that the BMO points lie within a plane (the BMO plane) that was used as a reference to spatially align each ONH on the same coordinate system.

### Digital Alignment of OCT volumes

To improve the performance of our deformation mapping protocol, it is necessary to remove rigid-body translations and rotations resulting from head and/or eye movements of the subjects in between OCT acquisitions. To this end, each OCT volume under a biomechanical load (adduction, abduction, or elevated IOP) was digitally aligned with its corresponding primary gaze OCT volume using a commercial software Amira (version 2020.1, FEI, Hillsboro, Oregon, USA), as described in our previous publication.^20^

### In Vivo Displacement and Strain Mapping of the ONH

We used a commercial digital volume correlation (DVC) module (Amira, 2020.3, Waltham, Massachussets: Thermo Fisher Scientific) to map the three-dimensional deformations between the baseline scan and scans under external loads (adduction, abduction, or elevated IOP) for each patient. This algorithm was used in our previous study to study associations between biomechanical strain and visual field loss.^20^ Briefly, each ONH morphology was sub-divided into ~4,000 cubic elements, and ~3,500 nodes (points), at which locations 3D displacements (vectors) were mapped following the application of external loads. We then derived the effective strain from the 3D displacements. The effective strain is a convenient local measure of 3D deformation that accounts for both compressive and tensile effects. In other words, the higher the compressive or tensile strain, the higher the effective strain.^34, 36, 37^ Details of the DVC algorithm used in this study, including how the strain values are derived and the validation of our DVC algorithm were provided in our previous paper.^20^

For each OCT volume, IOP- and gaze-induced effective strains were averaged across the entire ONH region within a 2,800 x 2,800 μm en-face region centered on the BMO center (**Figure 1**). To take into account potential variations in acute IOP elevation across eyes, the IOP-induced effective strains were normalized according to the following equation:

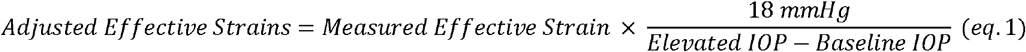

This approach was robust with respect to nonlinear stiffening of the sclera (validated in our previous work).^20^ It assumed linearity of the relationship, which was a reasonable assumption for IOP above 18 mmHg, according to the scleral strain response observed by Fazio et al.^38^ and Grytz et al.^39^ for human scleral strains, and by Midgett et al.^40^ for human LC strains.

### Statistical Analysis

Statistical analyses were performed using MATLAB (version 2018a, The MathWorks, Inc., Natick, Massachusetts, USA). Effective strains were defined as the continuous variable and the subjects’ diagnoses (control, HM, HMG, PM, and PM+S) were defined as categorical variables. To perform statistical analysis, we split our investigation into (1) differences in strains between HM and HMG under each load and (2) differences in strains between myopia spectrum (HM, PM and PM+S) under each load. We used independent samples t-test to compare the mean values of effective strain between the diagnostic groups. Analysis of variance (ANOVA) was used to determine the differences across groups in age, axial length, visual field MD, pattern standard deviation (PSD) and IOP on the day of the experiment. Linear regressions were used to study the associations between (1) axial length (or refractive error) and ONH effective strains; and those between (2) adduction-induced strains and IOP-induced strains. Statistical significance level was set at 0.05.

## Results

### Demographics

We excluded 6 HM subjects, 5 HMG subjects, 5 PM subjects and 11 PM+S from the study due to poor *en-face* LC visibility (<50% of the BMO area), and therefore 99 controls, 45 HM subjects, 30 HMG subjects, 16 PM and 64 PM+S subjects were included in the final analysis.

There were significant differences (p<0.05) in terms of age, axial length, visual field (MD and PSD values) between groups (**Table 1**). Notably, HM subjects were on average younger as compared to other groups. Myopic subjects (HM, HMG, PM and PM+S) exhibited a higher axial length as compared to the control group. HMG, PM and PM+S had poorer visual field indices (a more negative MD and a more positive PSD) as compared to the HM and control groups. There was no significant difference in presenting IOP on the day of experiment between groups.

**Table 1.**
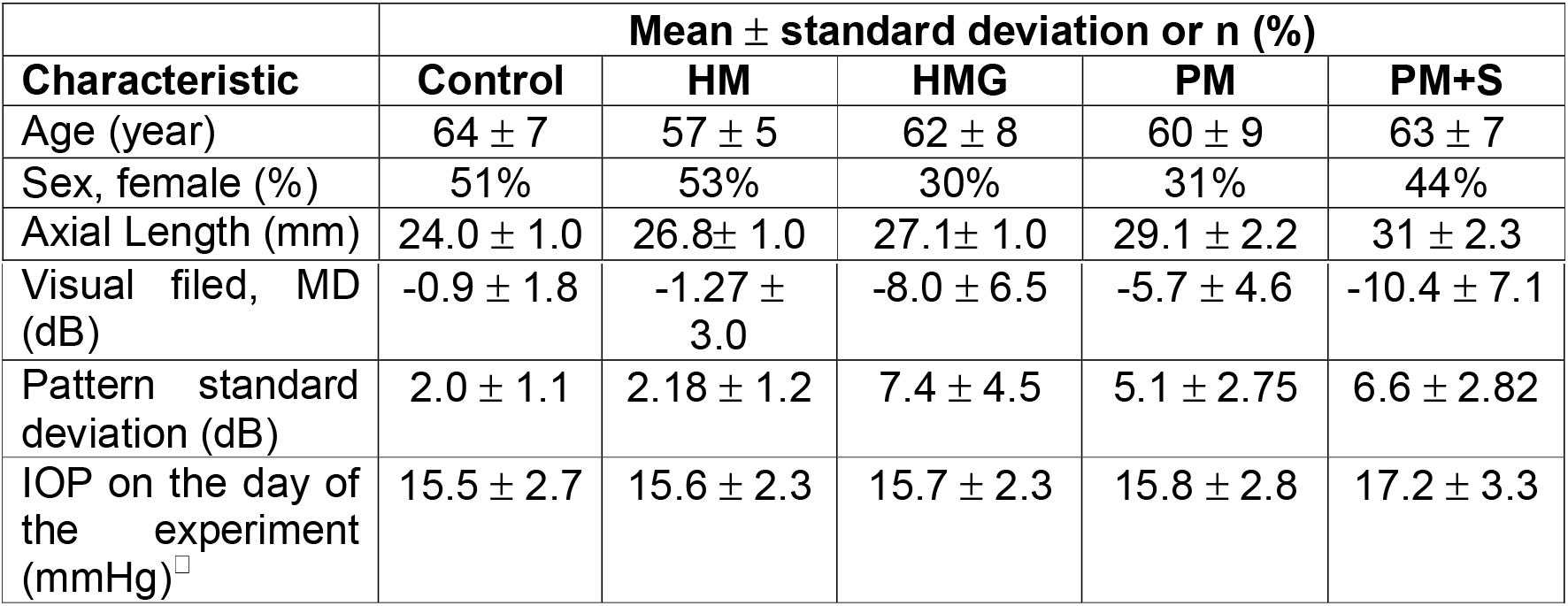
Characteristics of eyes in each group. ^This is IOP after treatment for HMG subjects.

| | Mean ± standard deviation or n (%) | | | | |
| --- | --- | --- | --- | --- | --- |
| Characteristic | Control | HM | HMG | PM | PM+S |
| Age (year) | 64 ± 7 | 57 ± 5 | 62 ± 8 | 60 ± 9 | 63 ± 7 |
| Sex, female (%) | 51% | 53% | 30% | 31% | 44% |
| Axial Length (mm) | 24.0 ± 1.0 | 26.8± 1.0 | 27.1± 1.0 | 29.1 ± 2.2 | 31 ± 2.3 |
| Visual field, MD (dB) | $-0.9 \pm 1.8$ | $-1.27 \pm 3.0$ | $-8.0 \pm 6.5$ | $-5.7 \pm 4.6$ | $-10.4 \pm 7.1$ |
| Pattern standard deviation (dB) | $2.0 \pm 1.1$ | $2.18 \pm 1.2$ | $7.4 \pm 4.5$ | $5.1 \pm 2.75$ | $6.6 \pm 2.82$ |
| IOP on the day of the experiment (mmHg) | $15.5 \pm 2.7$ | $15.6 \pm 2.3$ | $15.7 \pm 2.3$ | $15.8 \pm 2.8$ | $17.2 \pm 3.3$ |

### IOP-induced ONH strains

A visualization of strain maps (one eye from each diagnostic group) was given in **Figure 2**. Controls exhibited the least IOP-induced strains (3.6 ± 1.6%), followed by HM (3.9 ± 2.4%), HMG (4.7 ± 1.8%), PM (6.9 ± 5.0%) and PM+S (7.0 ± 5.2%).

**Figure 2.**
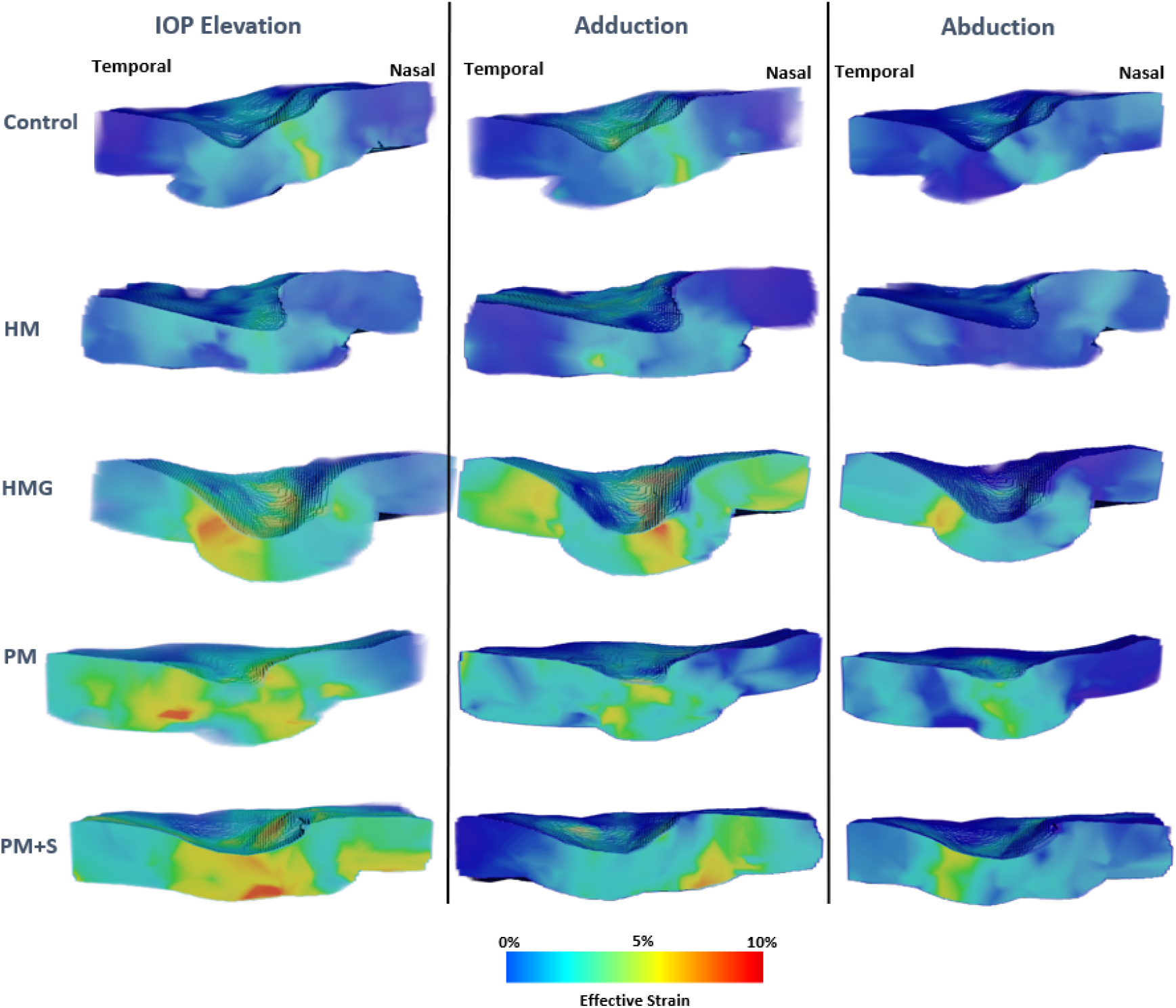
3-diensional effective strain maps of a sample eye (with central cut through the ONH) for each diagnostic group under loads.

We found that HM eyes exhibited significantly lower strains under IOP elevation than PM eyes (p < 0.001), HMG eyes (p = 0.04) and PM+S eyes (p < 0.001) (**Figure 3**). There was no significant difference in IOP-induced strains between control and HM eyes.

**Figure 3.**
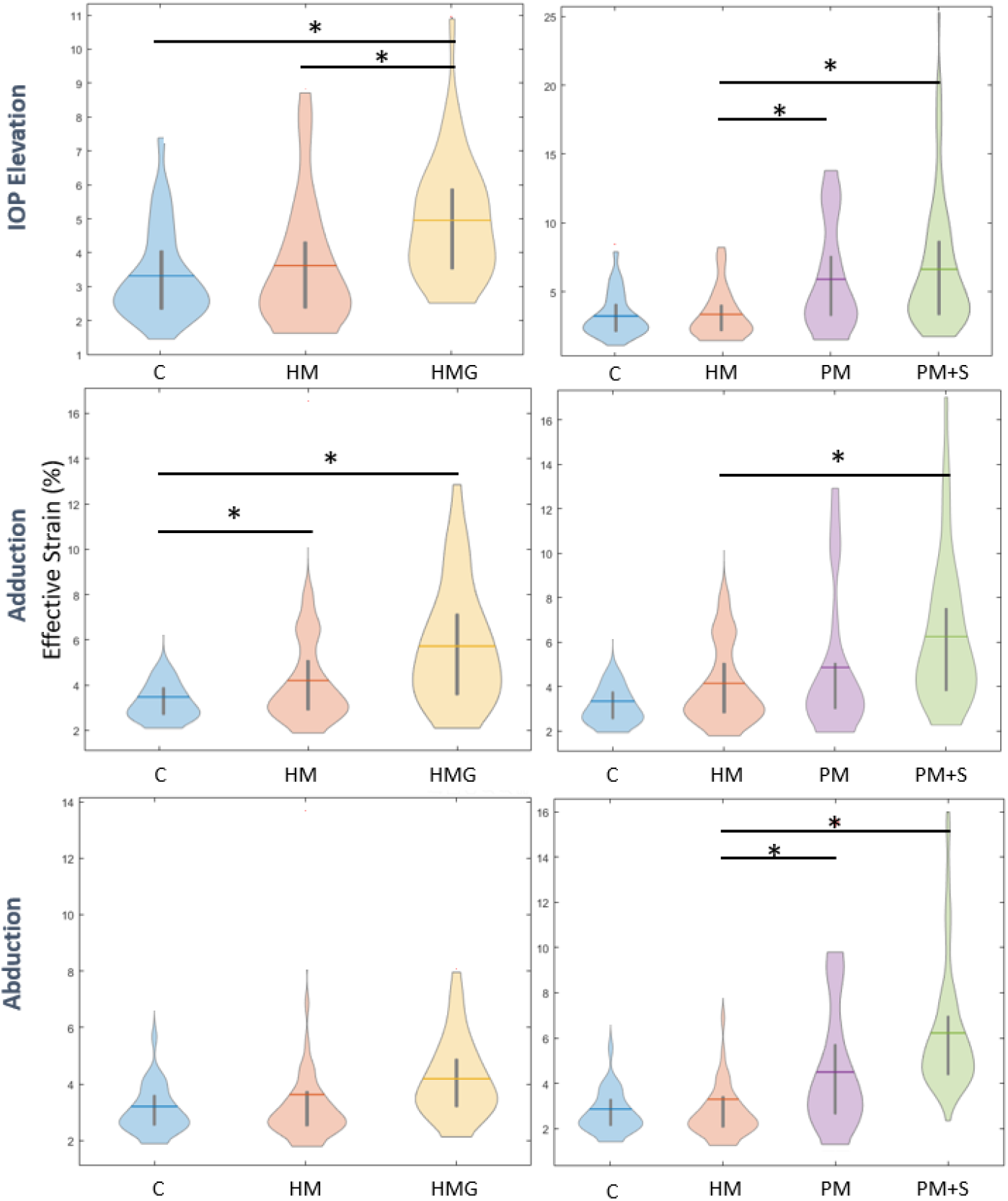
Violin-plot showing average ONH effective strains for each group under IOP elevation, Adduction and Abduction. Left column represents a comparison between HM and HMG and right column represents a comparison across myopia spectrum.* denotes significant differences between two groups.

### Gaze-induced ONH strains

Under abduction, we found a similar trend as that observed with IOP, with controls exhibiting the least strains (3.3 ± 1.4%), followed by HM (3.9 ± 2.4%), HMG (4.1 ± 1.4%), PM (5.0 ± 2.5%) and PM+S (5.3 ± 3.0%). Additionally, we found that PM+S eyes exhibited significantly higher strains than HM (p < 0.05) and PM eyes exhibited significantly higher strains than HM (p < 0.05) under 14 degrees abduction. We found no significant difference in abduction-induced strain between HM and controls.

Under adduction, we found that controls exhibited the least strains (3.8 ± 1.6%), followed by HM (4.8 ± 2.7%), PM (5.2 ± 2.5%), HMG (5.7 ± 2.5%), and PM+S (6.0 ± 3.1%). Additionally, we found that HM eyes exhibited significantly higher strain than controls (p < 0.05) and PM+S eyes exhibited significantly higher strain than HM eyes (p < 0.05) (**Figure 3**).

### Overall, Large ONH Strains (IOP-induced) Were Associated with High Axial Length or Low Refractive Error

Across all subjects, we found that the eyes that exhibited the largest IOP-induced ONH strains were also the eyes that exhibited the largest axial length (p<0.001, an axial length increase of approximately 2 mm results in an 1% increase in ONH strain) or the lowest refractive error (p<0.001, an worsening diopter of approximately 1 unit results in an 1% increase in ONH strain; **Figure 4**).

**Figure 4.**
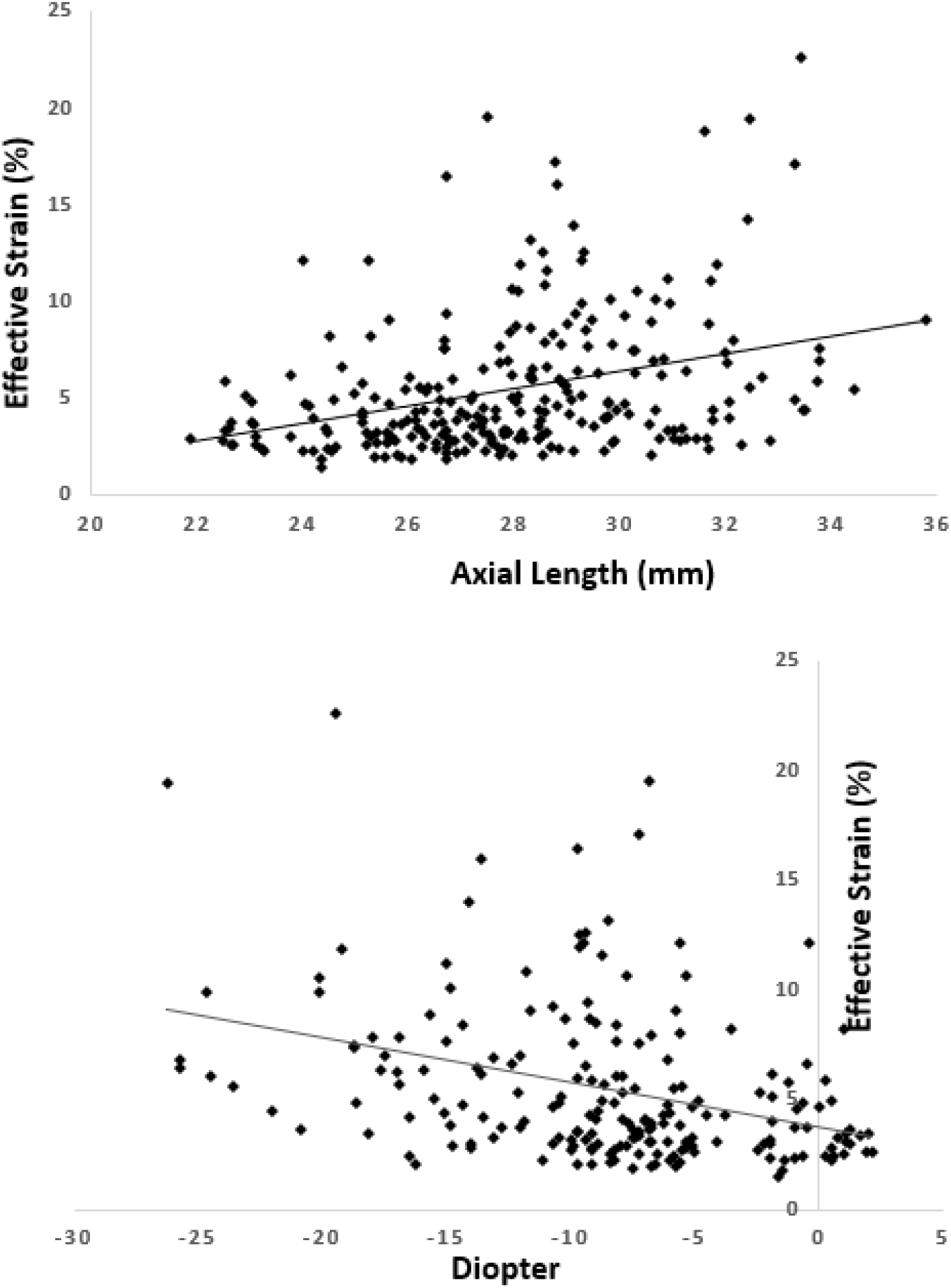
Scatterplot showing a significant linear relationship between ONH effective strain under IOP elevation and **(a)** axial length (mm) and **(b)** refractive error of all eyes.

### Adduction-Induced ONH Strains were Associated with IOP-Induced Strains

Across all eyes, we found that Adduction-Induced ONH Strains were positively associated with IOP-induced strains (p<0.001; **Figure 5**).

**Figure 5.**
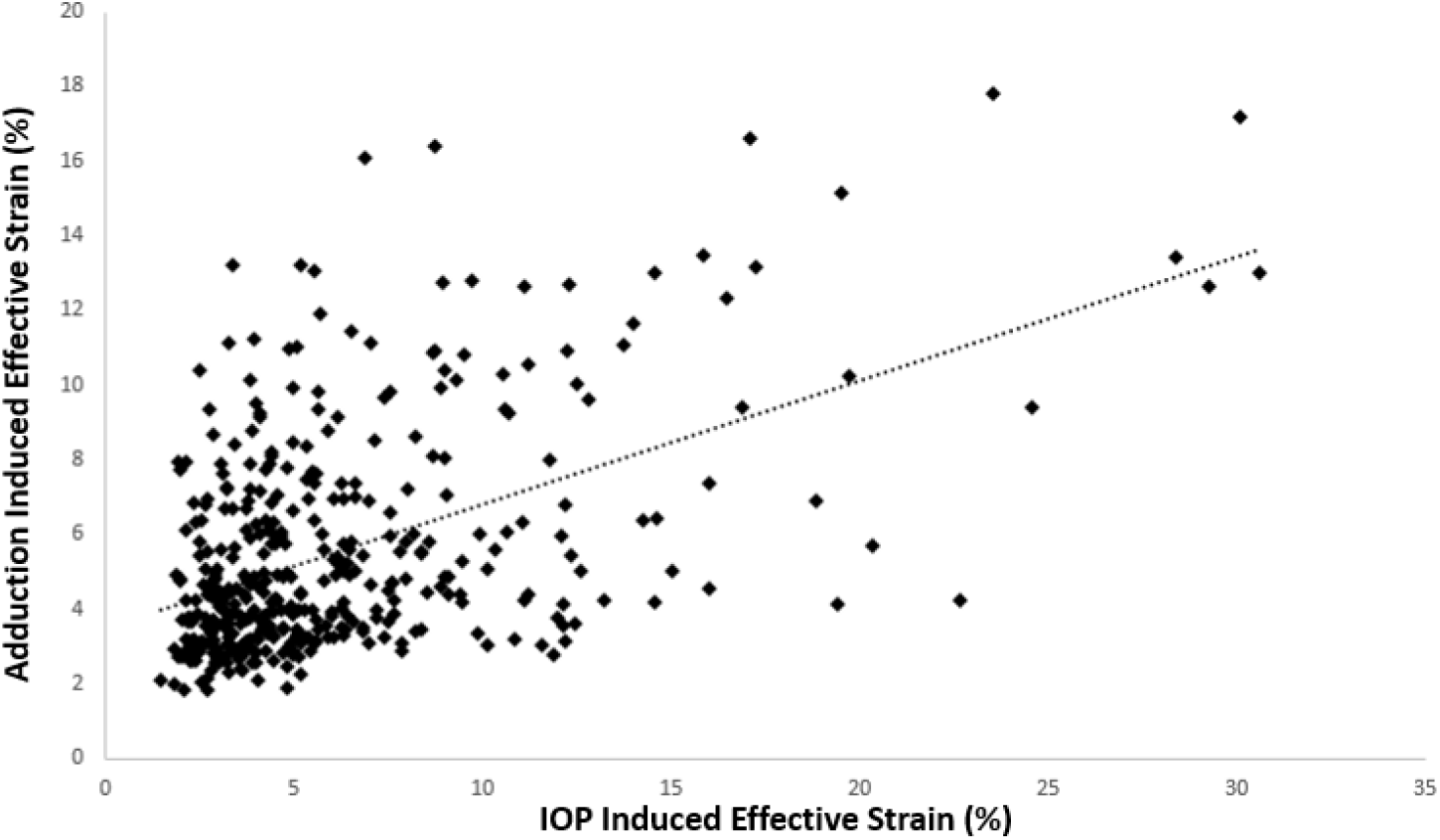
Scatterplot showing a significant positive linear relationship between ONH effective strain under IOP elevation and adduction.

## Discussion

In this study, we investigated the biomechanical responses of the ONH in eyes with ocular diseases (glaucoma, HM, and staphyloma) that have been suspected to have biomechanical origins. We found distinct differences in their biomechanical responses to two external loads (IOP and optic nerve traction). For IOP-induced strains, we found that control eyes exhibited the lowest strains, followed by HM, HMG, PM and PM+S. This trend was also observed in horizontal gaze and we found that IOP-induced strains and gaze-induced strains were correlated. Overall, the eyes with the ‘harshest’ conditions (PM+S) exhibited the highest strains under all conditions (7% under IOP elevation). In addition, our work suggests a significant differences in strain response (under acute IOP elevation) between HMG eyes and HM eyes, thus hinting at the possibility of using simple biomechanical tests to identify myopic eyes with underlying glaucoma.

In this study, we found that the ONH of HMG eyes exhibited higher levels of IOP-induced strain as compared to HM eyes. This may be explained by ONH excavation and deformation of the LC, a load bearing structure of the ONH, such as its thinning and posterior bowing in glaucoma.^41–45^ Jonas et al.^46^ found that among HM eyes, the presence of glaucoma was associated with thinner LC than in non-highly myopic eyes, which would promote a more structurally compliant response and thus greater strain.^47^ This is further supported by an ex vivo study by Midgett et al.,^48^ that found larger LC strain in more severely damaged glaucoma eyes compared to mildly damaged glaucoma eyes. They postulated that the increase in compliance was caused by ONH excavation, LC thinning and widening. Perhaps it is the compounded effect of myopia (scleral thinning^49,50^, tissue loss,^50^ failure at a lower load^49^ and increase in small diameter collagen fibrils^50^) and glaucoma (posterior laminar deformation^45,51^, scleral canal expansion^51,52^, posterior migration of anterior laminar insertion^53,54^, laminar thickness change^45,46^ and posterior bowing of the peripapillary sclera^55^) on ONH deformation and remodelling that results in greater strain in HMG eyes compared to HM eyes. Shifts in central retinal vessel trunks (CRVT) and branches during axial length elongation^56^ and during glaucoma development^57^ could be another reason behind the greater structural weakening of HMG eyes, since the CRVT (and branches) may provide mechanical strength to the ONH region.^58, 59^ Our result suggests that HMG eyes may have different biomechanical sensitivity as compared to HM eyes following the use of a simple biomechanical test. Future studies should aim to entangle the combined biomechanical influences of tissue remodelling associated to glaucoma and myopia development in terms of quantifiable strains, i.e., is a loss of ONH robustness caused by glaucoma remodeling greater than that caused by myopia associated remodelling? Additionally, we aim to employ similar methodology in a longitudinal cohort, following HM subjects who are at higher risks of developing glaucoma as a co-morbidity. This could help us understand the potential biomechanical link between the two conditions.

In this study, we also observed that PM eyes (especially those with staphylomas) exhibited significantly higher strains than HM eyes under all loads. Pathologic myopia (with the presence of staphylomas) is often associated with myopic traction maculopathy and both conditions are known to be responsible for structural weakening of the ocular globe.^60^ Specifically, posterior staphylomas are associated with thinning and elongation of the local sclera and deformity of the posterior ocular segment.^61^ Norman et al.^62^ showed that a thinner posterior sclera deformed more easily, and that scleral canal expansion and LC deformation would be larger than that seen with a thicker sclera. Thus, the presence of staphylomas could cause a significant structural weakening within the peripapillary sclera and our observation based on the in vivo strains supported this notion. For PM eyes without staphylomas, the finding of a marked increase in strains (all loads) was also reasonable, since PM is associated with a more extreme increase in axial length and further structural weakening of the sclera.^63^ We also suspect that a proportion of our PM subjects would go on to develop staphylomas, and we aim to monitor this closely.

In our work, we also found a strong association between axial length and ONH strains (all loads), and similarly between refractive error and ONH strains (all loads). It is well known from ex vivo studies that the sclera undergoes significant remodelling during myopia development,^13^ losing robustness (scleral weakening) in the process,^64^ resulting in scleral thinning (from the loss of collagen fibers and proteoglycans,^13, 64^ localized ectasia of the posterior sclera and eventually the elongation of the ocular globe (via a sliding of scleral lamellae as observed by an increase in sclera creep rate).^13, 64^ Additionally, the collagen fibers of the sclera had been observed to undergo re-alignment in myopic eyes, losing the protective circumferential alignment at the peripapillary region.^65^ It is also important to note that the sclera undergoes remodelling but not growth (resulting in an increase in surface area with no change in volume).^17, 66^ Thus, our findings indicated a distinct weakening of the sclera in a spectrum of myopia severity (defined by axial length). Axial length is also a risk factor in glaucoma^67, 68^ and the link between the two factors maybe a biomechanical one. A longitudinal study, which captures the ONH morphological changes and pathological changes in myopic group with a spectrum of axial length, would be of importance to strongly establish this relationship.

In this study, we found that the ONH of eyes with HM exhibited higher strain than that in controls eyes under adduction, but not under IOP elevation and abduction. Under adduction, the optic nerve sheath has been observed to exert significant tractional forces on the ONH which caused the ONH to deform in a distinct ‘see-saw’ pattern.^20, 22, 25, 69^ Our findings indicate that HM eyes were more susceptible to this tractional force as compared to controls, which was expected due to the weakening of the sclera during myopia development.^70^ Under IOP elevation, we found that both HM and controls exhibited the same level of strains – this was an intriguing finding, as we expected that the HM eyes would also be more sensitive to IOP elevation than controls. According to a review by Boote C. et al.,^71^ there were no studies which investigated the *in vivo* IOP-induced ONH strains between subject groups with different myopic status; thus, we cannot compare our findings directly with pre-existing results. Our non-significant finding under IOP elevation could be due to our relatively small number of subjects in the HM groups (51 subjects), and the fact that we used a binary division (25 mm in axial length) to separate non-HM from HM; several subjects in both groups had an axial length that were close to 25 mm. These subjects could affect the observed trends given our number of HM subjects and we plan to recruit more HM subjects to better elucidate this trend in our future study. It should be noted that our observation that HM eyes were more sensitive than control eyes was reasonable, as adduction had been shown to be associated with optic nerve traction,^24, 34, 72^ which could be a prevalent force that causes axial length elongation in myopia.^70^ Abduction was less associated with optic nerve traction and this may explain why we did not observe significant differences in strains between HM and controls.

To date, diagnosing glaucoma in myopic eyes is extremely challenging, due to the fact that structural biomarkers for glaucoma diagnosis are obscured by the structural alterations found in myopia. Here, we proposed simple biomechanical tests to map 3D ONH strains that could potentially be implemented clinically. We believe that a novel clinical marker to discriminate glaucoma (or its progression) in myopia would be of high importance. Our preliminary study has shown that this distinction may be possible with our biomechanical tests under IOP elevation and adduction. Given a strong correlation between IOP-induced strains and adduction induced strains, a non-invasive test using adduction to induce a biomechanical response is particularly attractive, as it requires no direct force application to the patient’s eye.

In this study, several limitations warrant further discussion. First, we could not define individual ONH tissues with clean boundaries in HM eyes; this was especially true for the PM eyes where boundary blurring was observed. This limited our ability to perform tissue-level analyses.

Second, OCT imaging could not capture the full thickness of the sclera in all eyes and thus our strains may not be representative of the posterior portion of the peripapillary sclera. Improvements in OCT hardware will be required to improve the vivo biomechanical measurements of the sclera.

Third, errors in both displacement and strain could occur. The errors observed here could arise from various sources such as OCT registration errors (intrinsic to the OCT machine), rotation of subjects’ head during OCT acquisition, OCT speckle noise and IOP fluctuations from ocular pulsations,^73^ all of which were difficult to control. However, displacement error magnitudes were still lower than the OCT voxel resolution;^20^ thus, these errors should not have significantly affected our observed trends.

Fourth, the use of ODM could introduce experimental errors as we applied an external force to the anterior sclera through the eyelid. We could not directly control the stress experienced at the ONH across subjects, and there may be variations due to the outflow facility^74^ and the biomechanical characteristics^75^ of each eye. Additionally, IOP also fluctuates during the scan duration due to the presence of the ocular pulse.^76^ The actual level of IOP raised via ODM could also potentially decline during the time needed for the OCT acquisition.^77^ Such a decline may also vary across subjects depending on the outflow facility^74^ and the biomechanical characteristics^75^ of each eye. Thus, this constituted another uncontrolled variable in our study.

Lastly, our HM subjects were relatively younger compared to controls; however, we believe that this would not significantly affect our result since our subjects were from an older population (more than 50 years of age) and notable changes in scleral elastin fibres and extracellular matrix should have occurred at an earlier age.^71^

In conclusion, this study provides in-vivo measurements of ONH strains in myopic eyes and investigated interactions between glaucoma and myopia. The simple mechanical test presented herein has the potential to be used to assess the level of mechanical strength of the myopic eye and to perhaps assist in the early detection of myopic eyes that will develop glaucoma and/or staphyloma.

## Abbreviations/Acronyms

ONH: Optic Nerve Head
IOP: Intraocular Pressure
LC: Lamina Cribrosa
HM: High Myopia
RNFL: Retinal Nerve Fiber Layer
PM: Pathologic Myopia
PM+S: Pathologic Myopia with Staphyloma
HMG: High Myopia with Glaucoma
OCT: Ocular Coherence Tomography
ODM: Ophthalmodynamometer
DVC: Digital Volume Correlation
CRVT: Central Retinal Vessel Trunks

## Acknowledgements

Acknowledgement is made to **(1)** the donors of the National Glaucoma Research, a program of the BrightFocus Foundation, for support of this research (G2021010S [MG]), **(2)** the NMRC-LCG grant ‘Tackling & Reducing Glaucoma Blindness with Emerging Technologies (TARGET)’, award ID: MOH-OFLCG21jun-0003 [MJAG] **(3)** the “Retinal Analytics through Machine learning aiding Physics (RAMP)” project supported by the National Research Foundation, Prime Minister’s Office, Singapore under its Intra-Create Thematic Grant “Intersection Of Engineering And Health” - NRF2019-THE002-0006 awarded to the Singapore MIT Alliance for Research and Technology (SMART) Centre, **(4)** SingHealth DukeNUS, Academic medicine research grant (AM/SU053/2021 [TAT]) **(5)** the National Natural Science Foundation of China (12002025 [XW]) and **(6)** the NMRC Clinician Scientist Award grant, award ID: MOH-CSAINV18may-0003 [QVH].

